# A meta-analysis of the diagnostic sensitivity and clinical utility of genome sequencing, exome sequencing and chromosomal microarray in children with suspected genetic diseases

**DOI:** 10.1101/255299

**Authors:** Michelle M. Clark, Zornitza Stark, Lauge Farnaes, Tiong Y. Tan, Susan M. White, David Dimmock, Stephen F. Kingsmore

**Affiliations:** Rady Children’s Institute for Genomic Medicine, San Diego, CA, USA; Murdoch Children’s Research Institute, Melbourne, Australia; Department of Pediatrics, University of California San Diego, San Diego, CA, USA; Department of Paediatrics, University of Melbourne, Melbourne, Australia.

**Author notes:** **Correspondence:** S Kingsmore. Rady Children’s Institute for Genomic Medicine, Rady Children’s Hospital, 3020 Children’s Way, San Diego, CA 92113, USA.

**Keywords:** clinical utility, diagnostic sensitivity, whole genome sequencing, whole exome sequencing, genetic disease, meta-analysis, chromosomal microarray, child, precision medicine, genomic medicine

## Abstract

**IMPORTANCE:** Genetic diseases are a leading cause of childhood mortality. Whole genome sequencing (WGS) and whole exome sequencing (WES) are relatively new methods for diagnosing genetic diseases.

**OBJECTIVES:** Compare the diagnostic sensitivity (rate of causative, pathogenic or likely pathogenic genotypes in known disease genes) and rate of clinical utility (proportion in whom medical or surgical management was changed by diagnosis) of WGS, WES, and chromosomal microarrays (CMA) in children with suspected genetic diseases.

**DATA SOURCES AND STUDY SELECTION:** Systematic review of the literature (January 2011 - August 2017) for studies of diagnostic sensitivity and/or clinical utility of WGS, WES, and/or CMA in children with suspected genetic diseases. 2% of identified studies met selection criteria.

**DATA EXTRACTION AND SYNTHESIS:** Two investigators extracted data independently following MOOSE/PRISMA guidelines.

**MAIN OUTCOMES AND MEASURES:** Pooled rates and 95% CI were estimated with a random-effects model. Metaanalysis of the rate of diagnosis was based on test type, family structure, and site of testing.

**RESULTS:** In 36 observational series and one randomized control trial, comprising 20,068 children, the diagnostic sensitivity of WGS (0.41, 95% CI 0.34-0.48, I^2^=44%) and WES (0.35, 95% CI 0.31-0.39, I^2^=85%) were qualitatively greater than CMA (0.10, 95% CI 0.08-0.12, I^2^=81%). Subgroup meta-analyses showed that the diagnostic sensitivity of WGS was significantly greater than CMA in studies published in 2017 (*P*<.0001, I^2^=13% and I^2^=40%, respectively), and the diagnostic sensitivity of WES was significantly greater than CMA in studies featuring within-cohort comparisons (*P*<001, I^2^=36%). Evidence for a significant difference in the diagnostic sensitivity of WGS and WES was lacking. In studies featuring within-cohort comparisons of singleton and trio WGS/WES, the likelihood of diagnosis was significantly greater for trios (odds ratio 2.04, 95% CI 1.62-2.56, I^2^=12%; *P*<.0001). The diagnostic sensitivity of WGS/WES with hospital-based interpretation (0.41, 95% CI 0.38-0.45, I^2^=50%) was qualitatively higher than that of reference laboratories (0.28, 95% CI 0.24-0.32, I^2^=81%); this difference was significant in meta-analysis of studies published in 2017 (*P*=.004, I^2^=34% and I^2^=26%, respectively). The rates of clinical utility of WGS (0.27, 95% CI 0.17-0.40, I^2^=54%) and WES (0.18, 95% CI 0.13-0.24, I^2^=77%) were higher than CMA (0.06, 95% CI 0.05-0.07, I^2^=42%); this difference was significant in meta-analysis of WGS vs CMA (*P*<.0001).

**CONCLUSIONS AND RELEVANCE:** In children with suspected genetic diseases, the diagnostic sensitivity and rate of clinical utility of WGS/WES were greater than CMA. Subgroups with higher WGS/WES diagnostic sensitivity were trios and those receiving hospital-based interpretation. WGS/WES should be considered a first-line genomic test for children with suspected genetic diseases.

**Key Points:** *Question:* What is the relative diagnostic sensitivity and clinical utility of different genome tests in children with suspected genetic diseases?

*Findings:* Whole genome sequencing had greater diagnostic sensitivity and clinical utility than chromosomal microarrays. Testing parent-child trios had greater diagnostic sensitivity than proband singletons. Hospital-based testing had greater diagnostic sensitivity than reference laboratories.

*Meaning:* Trio genomic sequencing is the most sensitive diagnostic test for children with suspected genetic diseases.

## Introduction

Genetic diseases (single gene disorders, genomic structural and chromosomal defects) are a leading cause of death in children less than ten years of age^1-8^. Establishing an etiologic diagnosis in children with suspected genetic diseases is important for timely implementation of precision medicine and optimal outcomes, particularly to guide weighty clinical decisions such as surgeries, extracorporeal membrane oxygenation, therapeutic selection, and palliative care^9^. With the exception of a few genetic diseases with pathognomonic findings at birth, such as chromosomal aneuploidies, etiologic diagnosis requires identification of the causative molecular basis. In practice, this is remarkably difficult for several reasons: Firstly, genetic heterogeneity - there are over 8,000 named single gene diseases^10^. Secondly, clinical heterogeneity - genetic disease presentations in infants are frequently *formes frustes* of classic descriptions in older children (see, for example Inoue et al.^11^). Thirdly, comorbidity is frequent in acutely ill infants – including prematurity, birth trauma, and sepsis – obfuscating clinical presentations^2^. Fourthly, approximately four percent of children have more than one genetic diagnosis^12^. Finally, disease progression is faster in children, switching the diagnostic odyssey to a race against time ^9,13,14^

Traditionally, establishment of molecular diagnoses was by serial testing guided by the differential diagnosis. CMA is the recommended first-line genomic test for children with several types of genetic diseases^15^. Serial testing employs many other tests – including newborn screening panels, metabolic testing, cytogenetics, chromosomal fluorescence *in situ* hybridization, single gene sequencing, and sequencing of panels of genes associated with specific disease types (such as sensorineural deafness, cardiac dysrhythmias, or epilepsy) ^15^. Iterative inquiry of differential diagnoses, however, frequently incurs a diagnostic odyssey and rarely allows etiologic diagnosis in time to influence acute management. Thus, inpatient management of children with suspected genetic diseases largely remains empiric, based on clinical diagnoses^9^.

Over the past five years, WGS and WES have started to gain broad use for etiologic diagnosis of infants and children with suspected genetic diseases^16-47^. By allowing concomitant examination of all or most genes in the differential diagnosis, WGS and WES have the potential to permit comprehensive and timely ascertainment of genetic diseases. Timely molecular diagnosis, in turn, has the potential to institute a new era of precision medicine for genetic diseases in children. During this period, WGS and WES methods have improved substantially. While numerous studies have been published^16-47^, there are not yet guidelines for their use by clinicians. Here we report a literature review and metaanalysis of the diagnostic sensitivity and rate of clinical utility of WGS and WES, compared with CMA, in children (age 0 to 18 years) with any suspected genetic disease.

## Methods

### Data Sources and Record Identification

We searched PubMed from January 1, 2011, to August 4, 2017 with the terms (“exome sequencing” or “whole genome sequencing” or “chromosomal microarray”), and (“diagnosis” or “clinical”), and “genetic disease” (Figure S1). We manually searched journals not indexed by PubMed that published articles related to clinical genomic testing. There were no language restrictions.

### Study Screening and Eligibility

Studies that evaluated the diagnostic sensitivity (proportion of patients tested who received genetic diagnoses) or clinical utility (proportion of patients tested in whom the diagnosis changed medical or surgical management) of WGS, WES and/or CMA were eligible. We limited eligibility to studies of cohorts with a broad range of genetic diseases, rather than one or a few disease types or clinical presentations, and in which the majority of probands were less than 18 years old. The systematic review and meta-analysis were performed according to the MOOSE and PRISMA guidelines (Table S1 and Figure S1).

### Inclusion Criteria and Data Extraction

Data extraction was manual. Data was reviewed for completeness and accuracy by at least two expert investigators and disparities were reconciled by consensus. The QUADAS-2 tool was used to assess the quality of the included studies (Table S2). The PICOTS typology of the criteria for inclusion of studies in quantitative analyses was:

#### Patients

Data extraction was limited to affected children (age less than 18 years) with suspected genetic disease.

#### Intervention

WGS, WES and/or CMA for etiologic diagnosis of a suspected genetic disease.

#### Comparator

The groups compared were subjects tested by WGS, WES and CMA. CMA was treated as the Reference Standard. Subgroups were patients tested as singletons (proband) and trios (parents and child).

#### Outcomes

Diagnostic sensitivity, rate of clinical utility. Molecular diagnoses were defined as pathogenic or likely pathogenic diplotypes affecting known or likely disease genes associated with phenotypes that overlapped at least part of the clinical features of the affected patient and that were reported to the patient’s clinician. Variants of uncertain significance and secondary findings were not extracted. The definition of clinical utility conformed to a position statement of the American College of Medical Genetics and Genomics, but was limited to changes in management for individual patients^63^.

#### Timing

Where more than one publication reported results from a cohort, we included the most recent value for diagnostic sensitivity. Clinical utility was assessed acutely (typically within six months of enrollment of the last patient).

##### Settings

Testing was performed clinically in hospital laboratories and reference laboratories, and experimentally in research laboratories.

##### Study Design

There were no study design restrictions.

#### Statistical Analysis

Between-study heterogeneity was explored by univariate analysis. Potential sources of heterogeneity included year of publication, number of probands, genetic disease tested, and consanguinity. The variable for genetic disease tested was treated as having four categories: Any genetic disease, neurodevelopmental and metabolic disorders, neurodevelopmental disabilities, and infants (average proband age less than one year at testing). The effect of disease tested on heterogeneity was explored with a random-effects model as described below. We used meta-regression to study associations of continuous variables (year, study size, and the rate of consanguinity) and heterogeneity.

When comparing rates between studies, raw proportions (i.e. molecular diagnostic and clinical utility rates) for individual studies were logit transformed due to small sample sizes and low event rates^48^. Pooled subgroup proportions and their variances were obtained by fitting an inverse-weighted logistic-normal random-effects model to the data. 95% confidence intervals (CIs) for individual studies were derived using the Clopper-Pearson exact method^49^. Pooled proportions and CIs were back-transformed for interpretation. For studies which conducted within-cohort comparisons, an inverse-weighted random-effects model was used to estimate pooled odds ratios (ORs). Due to the paired nature of the data, the marginal cross-over OR estimator of Becker and Balagtas^50,51^ was used for the meta-analysis of studies that conducted within-cohort comparisons of WES and CMA diagnostic rates. For all analyses, between-study heterogeneity was assessed using between-study variance (τ)^2^, the I^2^ statistic^52^ and Cochran’s Q test^53^. I^2^ values of 25%, 50%, and 75% indicate mild, moderate, and severe heterogeneity, respectively^52^. Subgroup analyses were conducted to minimize severe heterogeneity between studies. Subgroup differences in rates and ORs were tested when there was not significant evidence of within-group heterogeneity. Forest plots were used to summarize individual study and pooled group meta-analysis statistics. Two-tailed *P* ≤ .05 were considered statistically significant. All statistical analyses were conducted using the ‘meta’ (version 4.8.1) and ‘metafor’ (version 2.0.0) packages in R (version 3.3.3)^54-56^.

### Results

WGS and WES are relatively new methods for diagnosis of childhood genetic diseases. We compared the diagnostic sensitivity of WGS and WES with that of CMA, the recommended first-line genomic test for genetic diseases in children with intellectual disability, developmental delay, autism spectrum disorder and multiple congenital anomalies^15^. 2,093 records were identified by searches for studies of the diagnostic sensitivity of WGS, WES and CMA in affected children with a broad range of suspected genetic diseases (Figure S1). Thirty seven of these, featuring 20,068 children, met eligibility criteria and were included in qualitative analyses (Table S3 and S4)^16-47,57-62^. Thirty-six were case studies; one was a randomized controlled trial^25^. In these, the pooled diagnostic sensitivity of WGS was 0.41 (95% CI 0.34-0.48, seven studies, 374 children, I^2^=44%), which was qualitatively greater than WES (0.35, 95% CI 0.31-0.39, 26 studies, n=9,014, I^2^=85%) or CMA (0.10, 95% CI 0.08-0.12, 13 studies, n=11,429, I^2^=81%, Figure 1a). Severe heterogeneity (I^2^>75%) within the WES and CMA groups precluded statistical comparisons.

**Figure 1:**
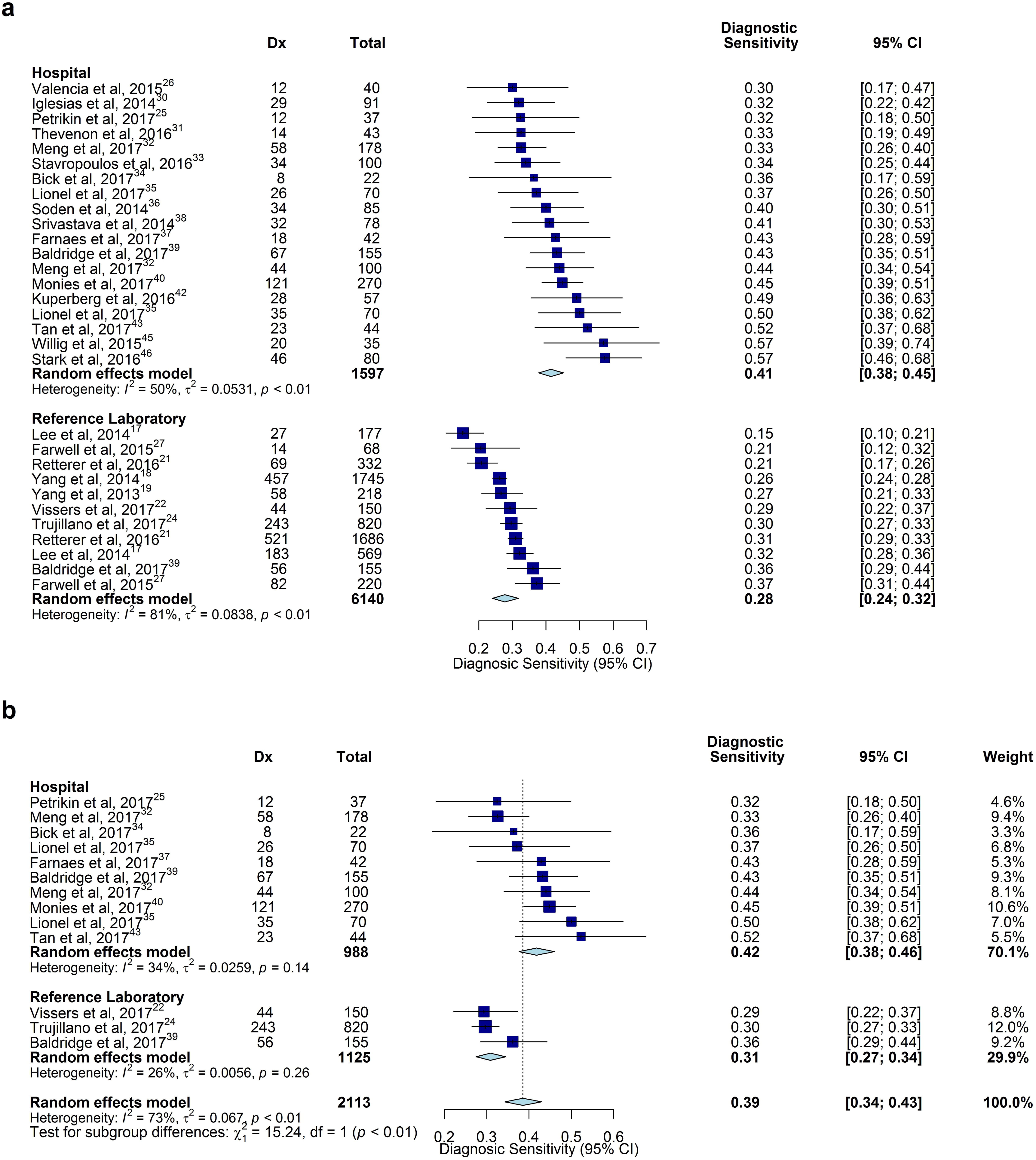
Comparison of diagnostic (Dx) sensitivity of WGS, WES and CMA. **a**. The pooled diagnostic sensitivity of WGS and WES were both greater than of CMA. However, severe heterogeneity precluded quantitative analysis, **b.** The subset of studies published in 2017 showed reduced heterogeneity for all subgroups. The pooled diagnostic sensitivity with WGS was significantly higher than with CMA (*P*<0001). **c.** Among manuscripts that provided complete data for the frequency of diagnoses made by WES and CMA, the pooled odds of diagnosis was 8.3 times greater for WGS (*P*<0001).

#### Analysis of heterogeneity of diagnostic sensitivity in studies of WGS and WES

We used meta-regression to model heterogeneity in the diagnostic sensitivity of WGS and WES. Studies varied in size from 22 to 1,745 probands; Meta-regression showed a modest relationship between study size and diagnostic sensitivity: On average, an increase of 1,000 subjects decreased the odds of diagnosis by 28% (Figure 2a, *P*=.01). Studies were published between 2013 and 2017; meta-regression showed that the odds of diagnosis increased by 14% each year (Figure 2b, *P*=.12). The rate of consanguinity varied between 0% and 100%. It was not significantly associated with the odds of diagnosis (*P*>.05). The proportion of diagnoses in which causal variants occurred *de novo* (rather than inherited) ranged from 0.18 - 0.70; meta-regression showed that a 10% increase in the rate of consanguinity decreased the odds of *de novo* variant diagnoses by 21% (*P*<.001; Figure 2c). Heterogeneity of diagnostic sensitivity in disease type and proband age subgroups precluded quantitative analysis (Figure S2).

**Figure 2:**
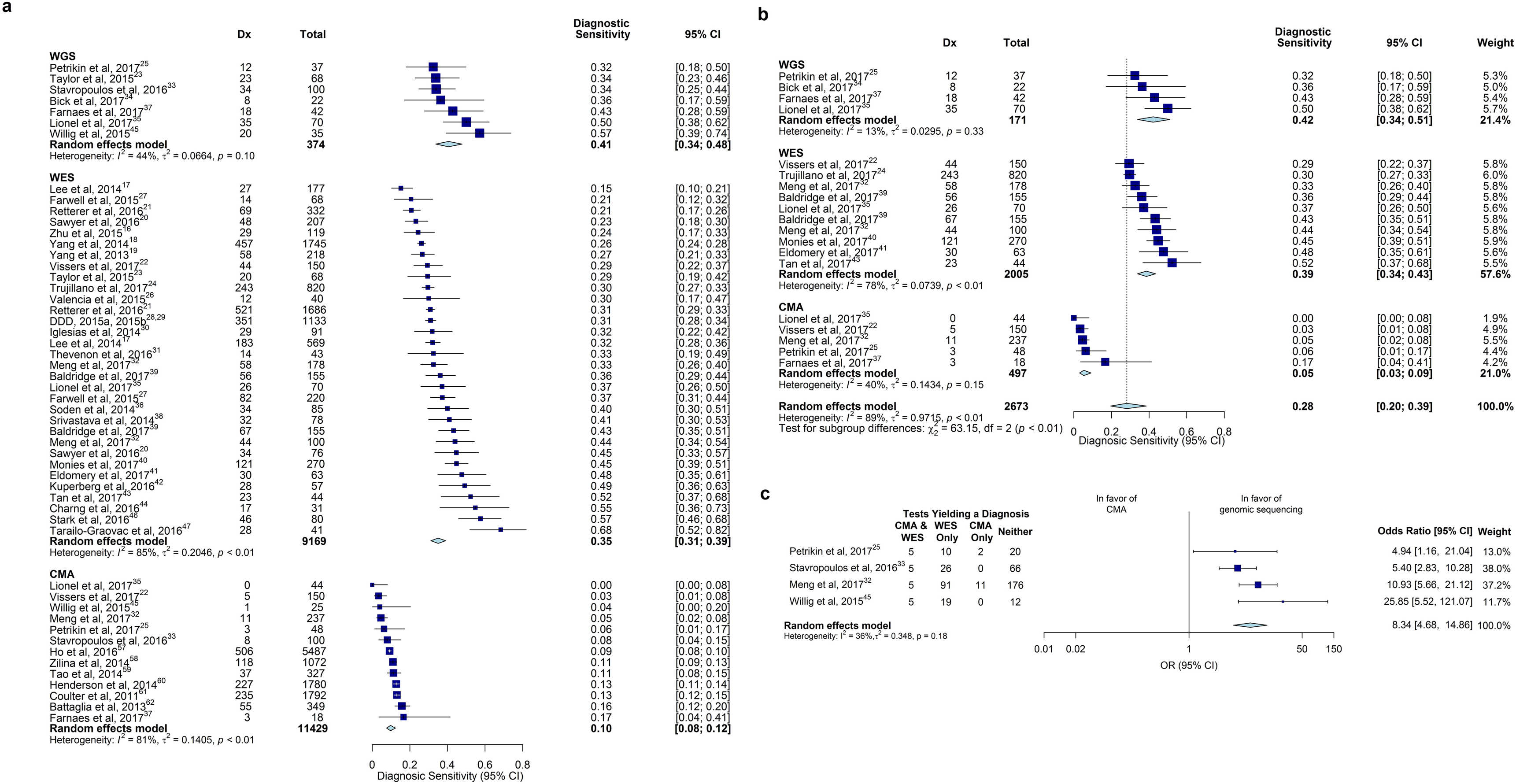
Exploration of heterogeneity of diagnostic sensitivity in WGS and WES studies. **a.** Meta regression scatterplot for study size. On average, an increase of 1000 subjects decreased the odds of diagnosis by 28% (*P*=.01). Size of data point corresponds to the study’s inverse-variance weight, **b.** Meta-regression scatterplot for year of study publication. On average, the odds of diagnosis increased by 14% per annum since 2013 (*P*=.12). **c.** The rate of diagnosis associated with *de novo* variation varied inversely with consanguinity. On average, increasing the rate of consanguinity by 10% decreased the odds of *de novo* variant diagnoses by 21% (*P*<.001).

#### Subgroup comparisons of diagnostic sensitivity of WGS, WES and CMA

Heterogeneity within WGS and CMA groups was mild following removal of variance associated with year of publication. In eleven studies of 1,962 children published in 2017, the pooled diagnostic sensitivity of WGS (0.42, 95% CI 0.34-0.51, I^2^=13%) was significantly greater than CMA (0.05, 95% CI 0.03-0.09, I^2^=40%; P<.0001,^Figure 1b^)^22,24,25,32,34,35,37,39,40,41,43^.

Only two studies, featuring 138 children, compared WES and WGS within cohorts. The diagnostic sensitivity of WES (0.29 and 0.37) did not differ significantly from that of WGS (0.34 and 0.50, respectively; *P*>.05)^23,35^. Since the diagnostic sensitivity of WES and WGS was not significantly different, we pooled WGS and WES studies in remaining subgroup analyses. Seven studies directly compared the proportion diagnosed by WGS or WES and CMA in 697 children; in each study, the diagnostic sensitivity of WGS/WES was at least three-fold higher than CMA^22,25,32,3,35,37,45^. Four of these manuscripts contained enough information to estimate the marginal odds ratios of receiving a diagnosis among subjects that received both WGS/WES and CMA^25,32,33,45^. In them, the odds of a diagnosis by WGS/WES was 8.3 times greater than CMA (95% CI, 4.7-14.9, I^2^=36%; *P*<.0001,Figure 1c).

#### Comparison of singleton and trio genomic sequencing and effect of site of testing

WGS/WES tests were either of affected probands or trios (proband, mother, father). In eighteen studies, comprising 3,935 probands, the heterogeneity of diagnostic sensitivity of singleton and trio WGS/WES was too great to permit quantitative analysis (Figure S3). Meta-analysis was performed in five studies (3,613 children) that compared the diagnostic sensitivity of WGS/WES by singleton and trio testing within cohorts^17,20,21,27,32^. In these studies, the odds of diagnosis using trios was double that using singletons (95% CI 1.62-2.56; I^2^=12%, *P*<.0001, Figure 3).

**Figure 3:**
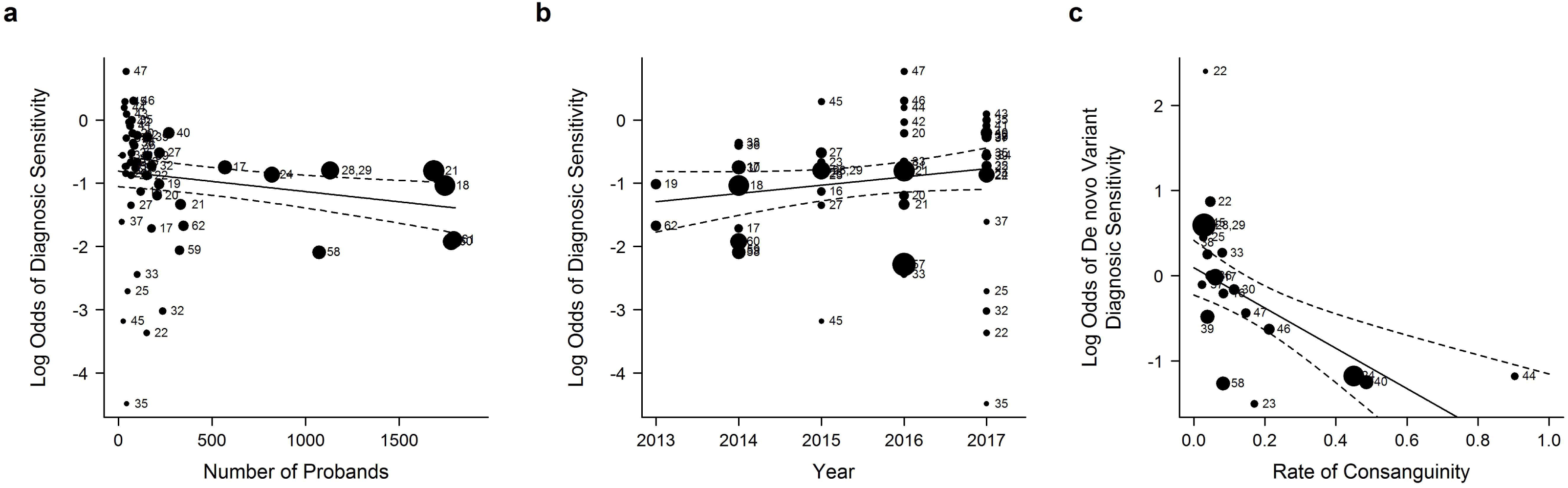
Comparison of diagnostic (Dx) sensitivity of singleton and trio WGS/WES in studies where both analyses were performed. In five studies that conducted within-cohort comparisons of singleton and trio genomic sequencing, the pooled odds of diagnosis for trios was twice that of singletons (*P*<.0001).

Studies were performed in three settings: i. Research studies of novel methods or disease gene discovery; ii. Clinical testing in hospital laboratories, where a deep phenotype was ascertained from the medical record at interpretation, and clinicopathologic correlation was facilitated by communication between clinicians and interpreters; iii. Reference laboratories, where phenotype information was limited to that provided in test orders, and communication between clinicians and interpreters was not possible. In nineteen studies, comprising 1,597 probands, the diagnostic sensitivity of hospital-based genomic sequencing was 0.41 (95% CI 0.38-0.45, I^2^=50%), and by reference laboratory-based genomic sequencing was 0.28 (95% CI 0.24-0.32, I^2^=81%, eleven studies, 6,140 probands, Figure 4a). Both hospital and reference laboratory subgroups demonstrated significant heterogeneity. However, heterogeneity was reduced in ten studies published in 2017 (I^2^=34%, *P*=.14, and I^2^=26%, *P*=.26, respectively)^22,24, 25,32,34,35,37,39,40,43^. In these, the diagnostic sensitivity of hospital genomic sequencing was 0.42 (95% CI 0.38-0.46, I^2^=34%), which was significantly higher than reference laboratories (0.31, 95% CI 0.27-0.34, I^2^=26%; *P=.004*, Figure 4b). Of note, hospital studies had an average of 84 subjects, while reference laboratory studies had an average of 558 subjects, providing a possible explanation for the inverse relationship between study size and rate of diagnosis (Figure 1a).

**Figure 4:**
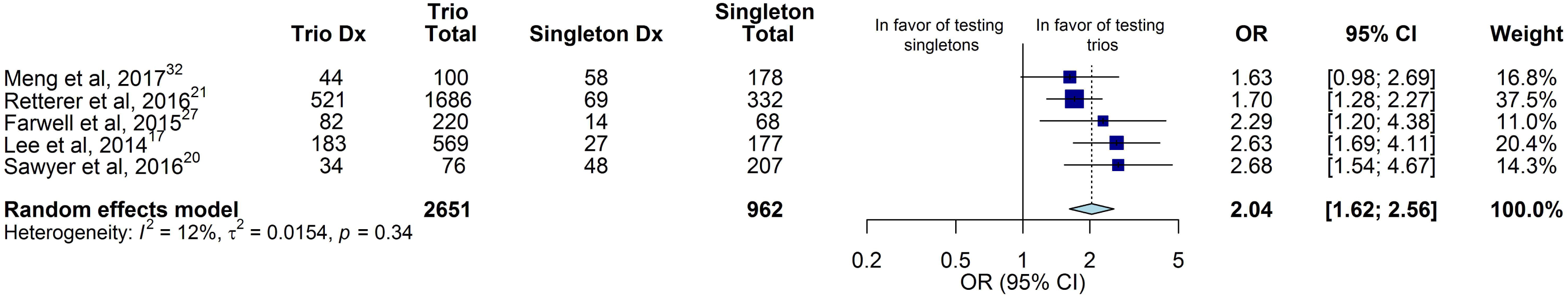
Comparison of diagnostic (Dx) sensitivity of WGS/WES in hospital laboratories and reference laboratories. **a.** The pooled diagnostic sensitivity of hospital-based testing was greater than reference laboratory testing. However, substantial heterogeneity was observed, **b.** The subset of studies published in 2017 showed reduced heterogeneity for both subgroups. The pooled diagnostic sensitivity was significantly greater in hospitals than in reference laboratories (*P*=.004).

#### Clinical Utility of WGS, WES and CMA

To decrease the heterogeneity in definitions of clinical utility between studies, we excluded cases in which the only change in clinical management was genetic counseling or reproductive planning^63^. The proportion of children receiving a change in clinical management by WGS results was 0.27 (95% CI 0.17-0.40, I^2^=54%, four studies of 136 children), compared with 0.18 (95% CI 0.13-0.24, I^2^=77%,twelve studies of 992 children) by WES, and 0.06 (95% CI 0.05-0.07, I^2^=42%, eight studies of 4,271 children) by CMA (Figure 5). Meta-analysis of WGS and CMA groups, for which heterogeneity was not significant (*P*=.09 and *P*=.10, respectively), demonstrated that the rate of clinical utility of WGS was higher than CMA (*P*<.0001)^25,32,34,35,37,45,59-61^. Meta-analysis of four studies that reported comparisons of rates of clinical utility by WGS/WES and CMA within cohorts^25,32,37,45^.

**Figure 5:**
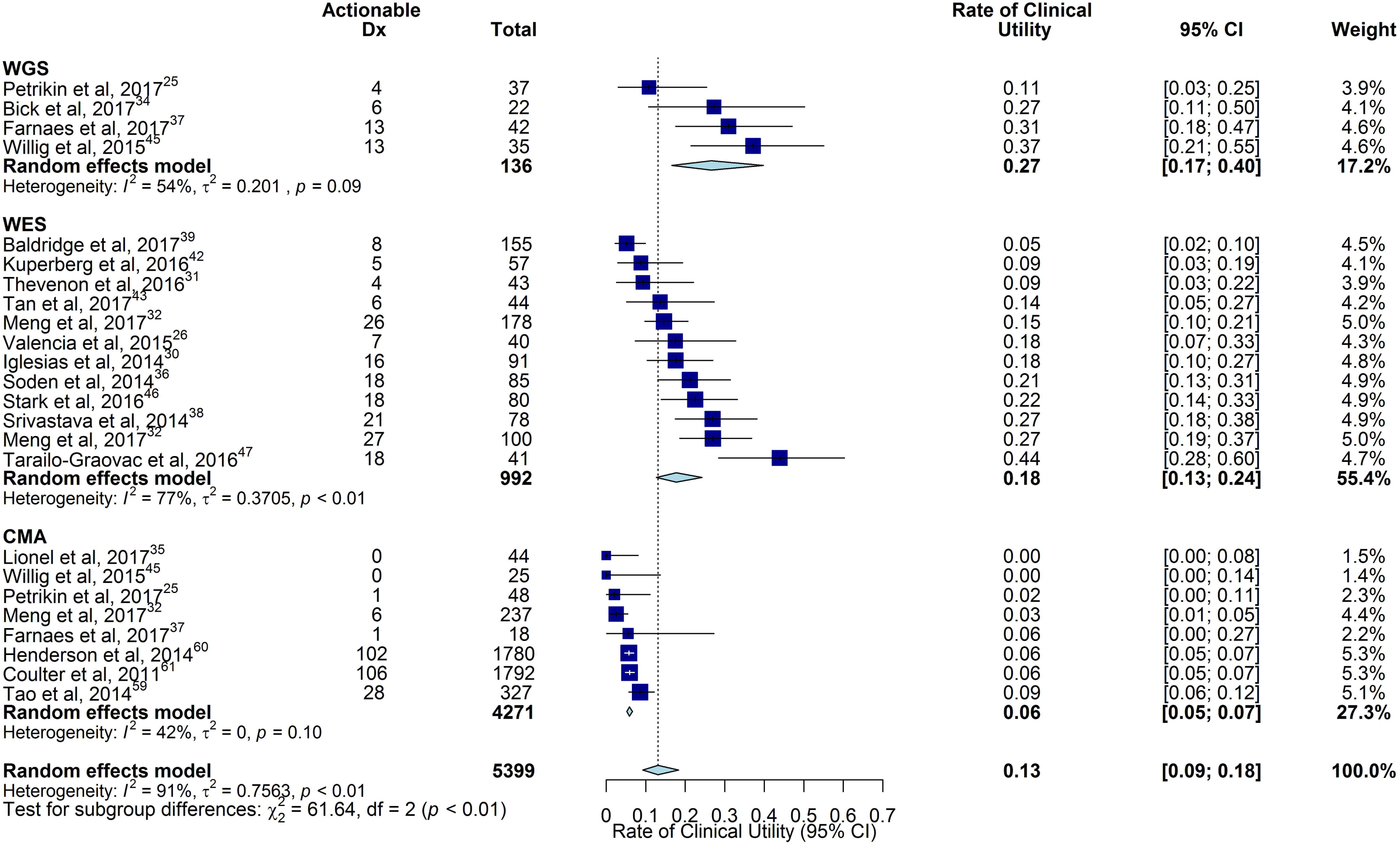
Comparison of the rate of clinical utility of WGS, WES and CMA. The rate of clinical utility was the proportion of children tested who received a change in medical or surgical management as a result of genetic disease diagnosis. The pooled rate of clinical utility of WGS and WES were both greater than of CMA. However, there was severe heterogeneity in the WES subgroup. Testing for subgroup differences amongst groups with low to moderate heterogeneity, we found that WGS diagnoses lead to an improved rate of clinical utility over CMA diagnoses.

### Discussion

Current guidelines state that CMA is the first-line genomic test for children with intellectual disability, developmental delay, autism spectrum disorder, and congenital anomalies^15,57-62,64-66^. Since 2011, WGS and WES have gained relatively broad use for etiologic diagnosis of genetic diseases, but guidelines do not yet exist for their use. A systematic review identified 37 publications in that period, comprising 20,068 affected children, which reported the diagnostic sensitivity of WGS, WES and/or CMA^16-47,57-62^. Since only thirteen (35%) of these reported results of a comparator test, pooling made comparisons susceptible to confounding from factors including clinical setting, patient factors, eligibility criteria, study quality, clinical expertise, and testing procedures. Meta-analysis of studies published in 2017, which removed variance associated with year of publication, showed that the diagnostic sensitivity of WGS (0.42, 95% CI 0.34-0.51, I^2^=13%) was significantly greater than CMA (0.05, 95% CI 0.03-0.09, I^2^=40%; *P*<.0001)^22,24,25,32,34,35,37,39,40,41,43^. Similarly, meta-analysis of studies featuring within-cohort comparisons showed that the odds of a diagnosis by WGS or WES was 8.3 times greater than CMA (95% CI, 4.7-14.9, I^2^=36%; *P*<.0001)^25,32,33,45^. These results suggest that CMA should no longer be considered the test with highest diagnostic sensitivity for childhood genetic diseases. Rather, WGS or WES should be considered a first-line genomic test for etiologic diagnosis of children with suspected genetic diseases.

While diagnostic sensitivity is an important measure of the value of a clinical test, the relative clinical utility of WGS, WES and CMA are more relevant for clinicians seeking to improve outcomes of rare childhood genetic diseases through implementation of precision medicine^9^. Given the genetic and clinical heterogeneity of genetic diseases^10^ and consequent myriad potential therapeutic interventions, it has been difficult to nominate meaningful, generally applicable measures of clinical utility. A previous approach was to collapse all interventions that were temporally and causally related to a molecular diagnosis into an overall “actionability” rate^25,35,37,45,59-61,63^ Such interventions were either based on practice guidelines endorsed by a professional society or peer-reviewed publications making medical management recommendations. While this has been applied in seven WGS and WES studies to date, definitions of actionability have varied. Furthermore, the evidence base for efficacy of ultra-rare genetic disease treatments is often qualitative rather than quantitative. Nevertheless, after excluding cases in which the only changes were ending the diagnostic odyssey or reproductive planning, WGS and WES had a higher actionability rates than CMA (0.27 [95% CI 0.17-0.40], 0.18 [95% CI 0.13-0.24], and 0.06 [95% CI 0.05-0.07], respectively). This difference was significant for WGS and CMA (*P*<.0001), in which within-group heterogeneity was not significant. One caveat was that children tested by CMA in these studies had multiple congenital anomalies, developmental delay, intellectual disability, or autism spectrum disorders, which were a subset of the presentations of children tested by WGS. Unfortunately no study has yet reported the relationship between clinical utility of WGS, WES or CMA and outcomes in children with genetic diseases.

Since WGS is about twice as expensive as WES, which is about twice as expensive as CMA, it is important to identify factors associated with high diagnostic sensitivity. One such factor was the test setting: Hospital laboratory testing had a higher diagnostic sensitivity (0.41, 95% CI 0.38-0.45) than reference laboratory testing (0.28, 95% CI 0.24-0.32). This difference was statistically significant (P=.004) among studies published in 2017, in which within-subgroup heterogeneity was not significant. This difference was supported by a study of double interpretation of WES of 115 children, first at a reference laboratory and second at the hospital caring for the children; the diagnostic sensitivity of reference laboratory interpretation was 0.33, and rate of false positive diagnoses was 0.03. The diagnostic sensitivity of hospital interpretation was 0.43, and there were no false positives^39^. The major difference between hospital and reference laboratory interpretation is the quality and quantity of phenotype information available at time of interpretation. In hospital testing, the phenotype is ascertained from the medical record, includes findings by subspecialist consultants, results of other concomitantly ordered tests, negative findings, and, in difficult cases, is supplemented by discussion with clinicians to ascertain material negative findings or clarify conflicting findings. In reference laboratories, the available phenotypes are those provided in test orders. They tend to be fewer in number and have less information content. One reference laboratory study found an association between the number of phenotypes available at interpretation and diagnostic yield: the diagnostic sensitivity was 0.26 with one to five phenotype terms, 0.33 with six to fifteen terms, and 0.39 with more than fifteen terms^24^. This was observed for all phenotypes, family structures, and inheritance patterns. Additional studies are needed to evaluate the reason for the apparent difference in diagnostic sensitivity of hospital and reference laboratory WES/WGS. In the interim, it is suggested that “send out” WES and WGS tests should be accompanied by as much phenotypic information as possible.

*De novo* variants accounted for the majority of genetic disease diagnoses, except in studies with high rates of consanguinity. Consanguinity is known to increase the population incidence of homozygous recessive genetic diseases. Herein, consanguinity was associated with decreased likelihood of attribution of diagnosis to *de novo* variants: Metaregression of 29 studies found the rate of consanguinity to be inversely related to the odds of diagnoses attributed to *de novo* variants (*P*<.001). Consanguinity is thought to increase the diagnostic sensitivity of WGS and WES: In one study, the diagnostic sensitivity of WES was 0.35 in 453 consanguineous families, and 0.27 in 443 non-consanguineous families^24^. However, meta-analysis failed to show a significant association between the rate of consanguinity and diagnostic sensitivity. Unfortunately, most studies did not report the proportion of probands with a family history of a similar illness, which was also anticipated to increase diagnostic sensitivity.

Testing of parent–child trios is considered superior to singleton (proband) testing for genetic disease diagnosis, since trios facilitate detection of *de novo* variants and allow phasing of compound heterozygous variants during interpretation. However, it is about twice as costly. Meta-analysis of five studies that compared the diagnostic sensitivity of singleton and trio testing within cohorts showed trio testing to have twice the odds of diagnosis than singleton testing (95% CI 1.62-2.56, *P*<.0001)^17,20,21,27,32^. This result was supported by a study in which 36% of unsolved singleton WES cases were diagnosed when re-analyzed as trios^18,19,41^. Additional studies are needed to guide clinicians with regard to the choice of initial trio or singleton testing. Factors to be considered include cost, time-to-result, and presence of consanguinity or family history of a similar condition.

Clinical WES has been much more broadly used than WGS, since WGS was very expensive until recently. WES examines almost all known exons and several hundred intronic nucleotides at ends of exons, or approximately two percent of the genome. WGS examines all exons and 90% of the genome. Only seven studies have reported the diagnostic sensitivity of clinical WGS in 374 children ^23,25,33-35,37,45^. Meta-analysis did not show the difference in the diagnostic sensitivity of WGS and WES to be significant. No study has yet directly compared the diagnostic sensitivity of WGS and WES. Additional studies are needed since the diagnostic sensitivity of WGS and WES are increasing disparately as a result of improved identification of disease-causing copy number and structural variations, repeat expansions, and non-exonic regulatory and splicing variations ^41,33,35,65-71^. In one recent study, these increased diagnostic sensitivity by 36%^41^. Recent research has shown WGS to have higher analytic sensitivity for copy number and structural variations than CMA, particularly small structural variations (less than 10,000 nucleotides ^33,35,71^), suggesting that WGS may become the single first-line genomic test for etiologic diagnosis of most children suspected to have a genetic disease. However, the published data do not yet support superiority of WGS over WES.

This meta-analysis had several limitations. Comparisons should be interpreted with caution due to heterogeneity of pooled averages of the published data. The highest level of evidence for clinical interventions is meta-analyses of randomized controlled trials (Level I)^72^. For WGS and WES, only one such study has yet been published. Published studies constitute Level II evidence (controlled studies or quasi-experimental studies) and Level III evidence (non-experimental descriptive studies, such as comparative studies, correlation studies, and case-control studies). The meta-analysis did not include diagnostic specificity (which has only been directly examined in one manuscript)^39^, nor the relative cost-effectiveness of WGS, WES and CMA, either in terms of the cost of the diagnostic odyssey or long term impact on healthcare utilization. It excluded next-generation sequencing-based panel tests, which are frequently used for specific presentations, such as epilepsy. It did not include subgroup analysis of the diagnostic sensitivity or clinical utility by affected organ system, which might have identified subgroups of children who are most likely to benefit from testing.

### Conclusions

In meta-analyses of 37 studies of children with suspected genetic diseases, WGS and WES had higher diagnostic sensitivity and rate of clinical utility than CMA, the current first-line genomic test for certain childhood genetic disorders. In a high proportion of children, diagnoses led to implementation of precision medicine. Additional randomized controlled studies are needed, particularly studies that examine the diagnostic determinants of optimal outcomes for children with rare genetic diseases^73^.

**Table 1.**
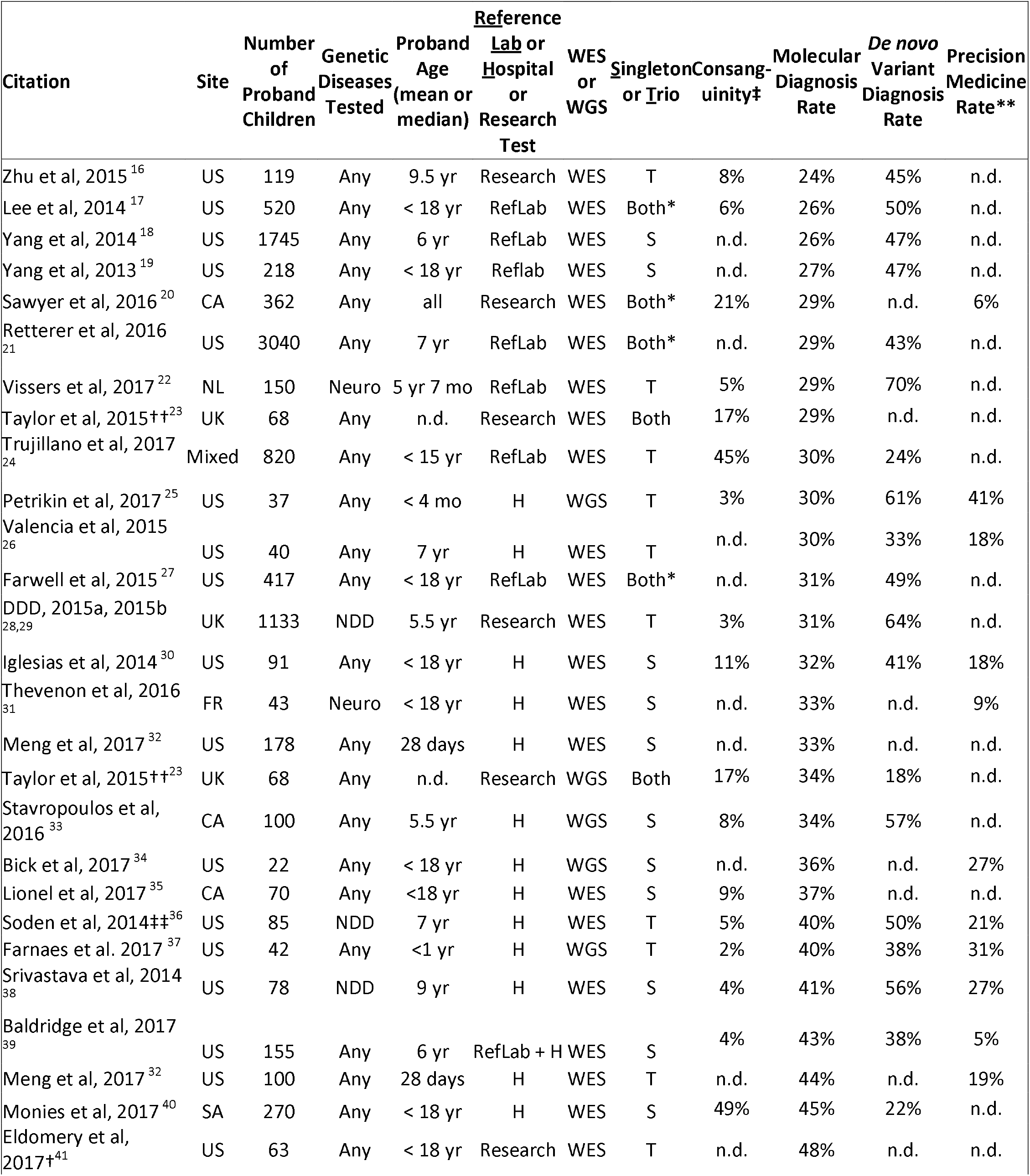
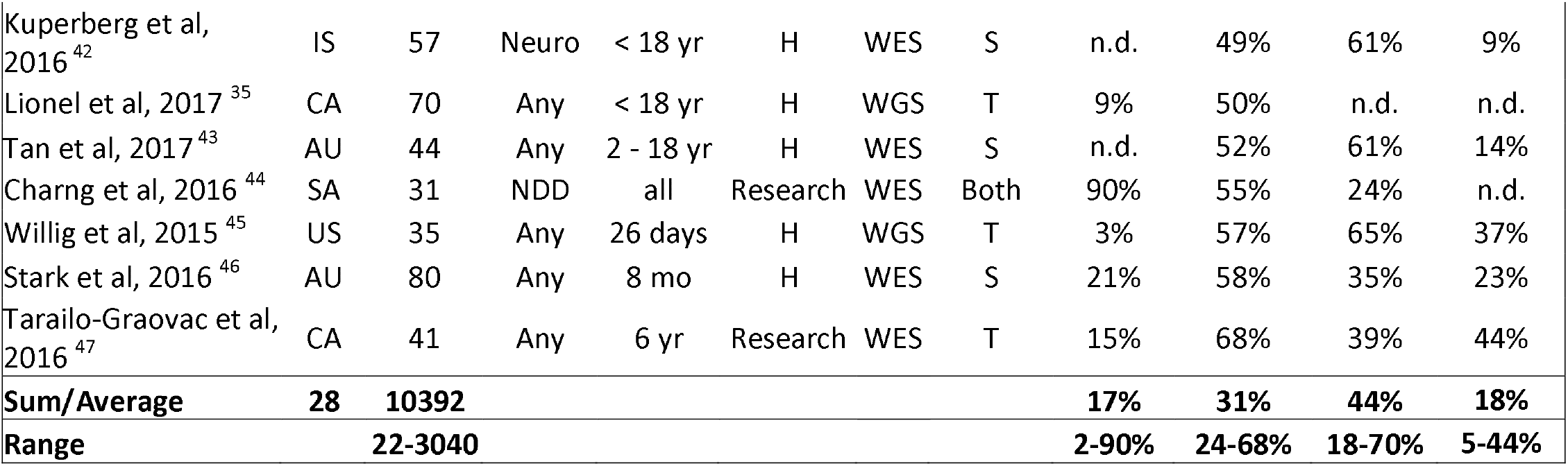
Characteristics of studies reporting diagnostic or clinical utility of whole exome sequencing or whole genome sequencing. ^*^Statistical difference between S and T within study. ^‡^By history or based on long runs of homozygosity. ^**^Other than reproductive plans. ^†^Unsolved by singleton WES. ^††^Bespoke methods for de novo variants. ^‡‡^Corrected to omit 15 infants reported in ref. 25. AU: Australia. CA: Canada. IS:Israel. NDD: neurodevelopmental disabilities. Neuro: neurologic. NL: Holland. SA: Saudi Arabia. UK: United Kingdom.

**Table 2.** Characteristics of studies reporting diagnostic or clinical utility of chromosomal microarray, whole exome sequencing or whole genome sequencing. ^*^Intellectual disability, developmental disorders, autism spectrum disorder, multiple congenital anomalies. ES: Estonia. IT: Italy. HK: Hong Kong. CA: Canada. NL: Holland.

| Citation | Site | Number of Proband Children | Genetic Diseases Tested | Proband Age (mean or median) | Consanguinity | Molecular Diagnosis Rate | *De novo* Variant Diagnosis Rate | Precision Medicine Rate |
| --- | --- | --- | --- | --- | --- | --- | --- | --- |
| Vissers et al, 2017 <sup>22</sup> | NL | 150 | Neuro | 5yr 7mo | 3% | 3% | 100% | n.d. |
| Meng et al, 2017 <sup>32</sup> | US | 278 | Any | 28 days | n.d. | 4% | n.d. | n.d. |
| Farnaes et al. 2017 <sup>37</sup> | US | 17 | Any | <1 yr | 6% | 6% | *n.d.* | 0% |
| Stavropoulos et al, 2016 <sup>33</sup> | CA | 100 | Neuro | 5.5 yr | n.d. | 8% | n.d. | n.d. |
| Ho et al, 2016 <sup>57</sup> | US | 5487 | * | 7.2 yr | n.d. | 9% | n.d. | n.d. |
| Zilina et al, 2014 <sup>58</sup> | ES | 1072 | Any | Postnatal | 8% | 11% | 22% | n.d. |
| Tao et al, 2014 <sup>59</sup> | HK | 327 | Any | <18 yr | n.d. | 11% | n.d. | 9% |
| Henderson et al, 2014 <sup>60</sup> | US | 1780 | * | <18 yr | n.d. | 13% | n.d. | 6% |
| Coulter et al, 2011 <sup>61</sup> | US | 1792 | * | < 18 yr | n.d. | 13% | n.d. | 6% |
| Battaglia et al, 2013 <sup>62</sup> | IT | 349 | * | <18 yr | n.d. | 16% | 45% | n.d. |
| **Sum/Average** | **9** | **11352** |  |  | **8%** | **11%** | **31%** | **6%** |

## ACKNOWLEDGEMENTS

We thank the many investigators who have built the evidence base for genomic medicine in children with genetic diseases. This work was supported by NICHD/NHGRI grant U19HD077693 to S.F.K. and by the Melbourne Genomics Health Alliance.

## References

1. March of Dimes Foundation Data Book for Policy Makers. Maternal, Infant, and Child Health in the United States 2016. http://www.marchofdimes.org/March-of-Dimes-2016-Databook.pdf Accessed 5/27/16

2. Xu J, Murphy SL, Kochanek KD, Arias E. Mortality in the United States, 2015. NCHS Data Brief. 2016 Dec;(267):1-8.

3. Wilkinson DJ, Fitzsimons JJ, Dargaville PA, et al. Death in the neonatal intensive care unit: changing patterns of end of life care over two decades. Arch Dis Child Fetal Neonatal Ed 2006; 91: F268–71.

4. Hagen CM, Hansen TW. Deaths in a neonatal intensive care unit: a 10-year perspective. Pediatr Crit Care Med 2004; 5: 463–68.

5. Ray JG, Urquia ML, Berger H, Vermeulen MJ. Maternal and neonatal separation and mortality associated with concurrent admissions to intensive care units. CMAJ 2012; 184: E956–62.

6. Yoon PW, Olney RS, Khoury MJ, Sappenfield WM, Chavez GF, Taylor D. Contribution of birth defects and genetic diseases to pediatric hospitalizations. A population-based study. Arch Pediatr Adolesc Med 1997; 151:1096–103.

7. O’Malley M, Hutcheon RG. Genetic disorders and congenital malformations in pediatric long-term care. J Am Med Dir Assoc 2007; 8: 332–34.

8. Stevenson DA, Carey JC. Contribution of malformations and genetic disorders to mortality in a children’s hospital. Am J Med Genet A 2004; 126A: 393–97.

9. Petrikin JE, Willig LK, Smith LD, Kingsmore SF. Rapid whole genome sequencing and precision neonatology. Semin Perinatol. 2015 Dec;39(8):623-31.

10. http://www.omim.ore/statistics/entry Accessed 5/27/2017.

11. Inoue S, Mangat C, Rafe’e Y, Sharman M. Forme Fruste of HLH (haemophagocytic lymphohistiocytosis): diagnostic and therapeutic challenges. BMJ Case Rep. 2015 Jan 29;2015.

12. Posey JE, et al. Resolution of Disease Phenotypes Resulting from Multilocus Genomic Variation. N Engl J Med. 2016 Dec 7.

13. Saunders CJ, et al. Rapid whole-genome sequencing for genetic disease diagnosis in neonatal intensive care units. Sci Transl Med. 2012 Oct 3;4(154):154ra135.

14. Miller NA, et al. A 26-hour system of highly sensitive whole genome sequencing for emergency management of genetic diseases. Genome Med. 2015 Sep 30;7(1):100.

15. Miller DT, et al. Consensus statement: chromosomal microarray is a first-tier clinical diagnostic test for individuals with developmental disabilities or congenital anomalies. Am J Hum Genet. 2010 May 14;86(5):749-64.

16. Zhu X, Petrovski S, Xie P, et al. Whole-exome sequencing in undiagnosed genetic diseases: interpreting 119 trios. Genet Med. 2015 Oct;17(10):774-81.

17. Lee H, Deignan JL, Dorrani N, et al. Clinical exome sequencing for genetic identification of rare Mendelian disorders. JAMA. 2014 Nov 12;312(18):1880-7.

18. Yang Y, Muzny DM, Xia F, et al. Molecular findings among patients referred for clinical whole-exome sequencing. JAMA. 2014 Nov 12;312(18):1870-9.

19. Yang Y, Muzny DM, Reid JG, et al. Clinical whole-exome sequencing for the diagnosis of mendelian disorders. N Engl J Med. 2013 Oct 17;369(16):1502-11.

20. Sawyer SL, Hartley T, Dyment DA, et al. Utility of whole-exome sequencing for those near the end of the diagnostic odyssey: time to address gaps in care. Clin Genet. 2016 Mar;89(3):275-84.

21. Retterer K, et al. Clinical application of whole-exome sequencing across clinical indications. Genet Med. 2016 Jul;18(7):696-704.

22. Vissers LE, et al. A clinical utility study of exome sequencing versus conventional genetic testing in pediatric neurology. Genet Med. 2017 Mar 23.

23. Taylor JC, et al. Factors influencing success of clinical genome sequencing across a broad spectrum of disorders. Nat Genet. 2015 47(7):717-26.

24. Trujillano D, et al. Clinical exome sequencing: results from 2819 samples reflecting 1000 families. Eur J Hum Genet. 2017 Feb;25(2):176-182.

25. Petrikin JE, Cakici JA, Clark MM, et al. Rapid Whole Genome Sequencing for Etiologic Diagnosis of Critically Ill Infants: The NSIGHT1 Randomized Controlled Trial. bioRxiv 218255; doi: https://doi.org/10.1101/218255

26. Valencia CA, et al. Clinical Impact and Cost-Effectiveness of Whole Exome Sequencing as a Diagnostic Tool: A Pediatric Center’s Experience. Front Pediatr. 2015 3:67.

27. Farwell KD, et al. Enhanced utility of family-centered diagnostic exome sequencing with inheritance model-based analysis: results from 500 unselected families with undiagnosed genetic conditions. Genet Med. 2015 Jul;17(7):578-86.

28. Deciphering Developmental Disorders Study. Large-scale discovery of novel genetic causes of developmental disorders. Nature. 201 519(7542):223-8.

29. Wright CF, et al. Genetic diagnosis of developmental disorders in the DDD study: a scalable analysis of genome-wide research data. Lancet. 2015 385(9975):1305-14.

30. Iglesias A, Anyane-Yeboa K, Wynn J, et al. The usefulness of whole-exome sequencing in routine clinical practice. Genet Med. 2014 16(12):922-31.

31. Thevenon J, Duffourd Y, Masurel-Paulet A, et al. Diagnostic odyssey in severe neurodevelopmental disorders: toward clinical whole-exome sequencing as a first-line diagnostic test. Clin Genet. 2016 89(6):700-7.

32. Meng L, Pammi M, Saronwala A, et al. Utility of Exome Sequencing for Infants in Intensive Care Units: Ascertainment of Severe Single-Gene Disorders and Impact on Medical Management. JAMA Pediatr. 2017 Dec 4;171(12):e173438.

33. Stavropoulos DJ, et al. Whole-genome sequencing expands diagnostic utility and improves clinical management in paediatric medicine, npj Genomic Med 2016 1:15012.

34. Bick D, Fraser PC, Gutzeit MF, et al. Successful Application of Whole Genome Sequencing in a Medical Genetics Clinic. J Pediatr Genet. 2017 6(2):61-76.

35. Lionel AC, Costain G, Monfared N, et al. Improved diagnostic yield compared with targeted gene sequencing panels suggests a role for whole-genome sequencing as a first-tier genetic test. Genet Med. 2017. August 3.

36. Soden SE, Saunders CJ, Willig LK, et al. Effectiveness of exome and genome sequencing guided by acuity of illness for diagnosis of neurodevelopmental disorders. Sci Transl Med. 2014 6(265):265ra168.

37. Farnaes L, Amber Hildreth A, Nathaly M. Sweeney NM, et al. Precision Medicine through Rapid Whole Genome Sequencing Decreases Morbidity and Healthcare Cost of Inpatient Infants. bioRxiv 253534; doi: https://doi.org/10.1101/253534

38. Srivastava S, Cohen JS, Vernon H, et al. Clinical whole exome sequencing in child neurology practice. Ann Neurol. 2014 76(4):473-83.

39. Baldridge D, et al. The Exome Clinic and the role of medical genetics expertise in the interpretation of exome sequencing results. Genet Med. 2017 Mar 2.

40. Monies D, Abouelhoda M, AlSayed M, et al. The landscape of genetic diseases in Saudi Arabia based on the first 1000 diagnostic panels and exomes. Hum Genet. 2017 136(8):921-939.

41. Eldomery MK, et al. Lessons learned from additional research analyses of unsolved clinical exome cases. Genome Med. 2017 Mar 21;9(1):26.

42. Kuperberg M, Lev D, Blumkin L, et al. Utility of Whole Exome Sequencing for Genetic Diagnosis of Previously Undiagnosed Pediatric Neurology Patients. J Child Neurol. 2016 31(14):1534-1539.

43. Tan TY, Dillon OJ, Stark Z, et al. Diagnostic Impact and Cost-effectiveness of Whole-Exome Sequencing for Ambulant Children With Suspected Monogenic Conditions. JAMA Pediatr. 2017 Jul 31.

44. Charng WL, Karaca E, Coban, et al. Exome sequencing in mostly consanguineous Arab families with neurologic disease provides a high potential molecular diagnosis rate. BMC Med Genomics. 2016 9(1):42

45. Willig LK, et al. Whole-genome sequencing for identification of Mendelian disorders in critically ill infants: a retrospective analysis of diagnostic and clinical findings. Lancet Respir Med. 2015 May;3(5):377-87.

46. Stark Z, et al. A prospective evaluation of whole-exome sequencing as a first-tier molecular test in infants with suspected monogenic disorders. Genet Med. 2016 Nov;18(11):1090-1096.

47. Tarailo-Graovac M, et al. Exome Sequencing and the Management of Neurometabolic Disorders. N Engl J Med. 2016 Jun 9;374(23):2246-55.

48. Hamza TH, van Houwelingen HC, Stijnen T. The binomial distribution of meta-analysis was preferred to model within-study variability. J Clin Epidemiol. 2008, 61:41–51.

49. Clopper CJ, Pearson ES. The use of confidence or fiducial limits illustrated in the case of the binomial. Biometrika. 1934, 26(4):404–413.

50. Becker MP, Balagtas CC. Marginal modeling of binary cross-over data. Biometrics. 1993, 49: 997-1009.

51. Curtin F, Elbourne D, Altman DG. Meta-analysis combining parallel and cross-over clinical trials. II: Binary outcomes. Statistics in Medicine. 2002, 21: 2145-2159.

52. Higgins J, Thompson SG. Quantifying heterogeneity in a meta-analysis. Statistics in medicine. 2002, 21:1539–1558.

53. Cochran WG. The Comparison of Percentages in Matched Samples. Biometrika. 1950, 37(3/4): 256-266.

54. R Core Team. R: A language and environment for statistical computing. R Foundation for Statistical Computing, Vienna, Austria. 2016. URL https://www.R-proiect.org/.

55. Viechtbauer, W. Conducting meta-analyses in R with the metafor package. Journal of Statistical Software. 2010 36(3), 1–48. http://www.istatsoft.ore/v36/i03/

56. Schwarzer G. meta: An R package for meta-analysis. R News. 2007, 7(3): 40–5.

57. Ho KS, et al. Clinical Performance of an Ultrahigh Resolution Chromosomal Microarray Optimized for Neurodevelopmental Disorders. Biomed Res Int. 2016:3284534.

58. Zilina O, et al. Chromosomal microarray analysis as a first-tier clinical diagnostic test: Estonian experience. Mol Genet Genomic Med. 2014 Mar;2(2):166-75.

59. Tao VQ, Chan KYK, Chu YWY, et al. The Clinical Impact of Chromosomal Microarray on Paediatric Care in Hong Kong. PLoS ONE 2014 9(10): e109629.

60. Henderson LB, Applegate CD, Wohler E, Sheridan MB, Hoover-Fong J, Batista DA. The impact of chromosomal microarray on clinical management: a retrospective analysis. Genet Med. 2014 Sep;16(9):657–64.

61. Coulter ME, Miller DT, Harris DJ, et al. (2011) Chromosomal microarray testing influences medical management. Genet Med 13: 770-776.

62. Battaglia A, et al. Confirmation of chromosomal microarray as a first-tier clinical diagnostic test for individuals with developmental delay, intellectual disability, autism spectrum disorders and dysmorphic features. Eur J Paediatr Neurol. 2013 Nov;17(6):589-99.

63. ACMG Board of Directors. Clinical utility of genetic and genomic services: a position statement of the American College of Medical Genetics and Genomics. Genet Med. 2015;17(6):505-7.

64. Tammimies K, et al. Molecular Diagnostic Yield of Chromosomal Microarray Analysis and Whole-Exome Sequencing in Children With Autism Spectrum Disorder. JAMA. 2015;314(9):895-903.

65. Manning M, Hudgins L, et al. Array-based technology and recommendations for utilization in medical genetics practice for detection of chromosomal abnormalities. Genet Med. 2010;12(11):742-745.

66. Moeschler JB, Shevell M. Comprehensive evaluation of the child with intellectual disability or global developmental delays. Pediatrics. 2014; 134(3):e903-e918.

67. Känsäkoski J, Jääskeläinen J, Jääskeläinen T, et al. Complete androgen insensitivity syndrome caused by a deep intronic pseudoexon-activating mutation in the androgen receptor gene. Sci Rep. 2016 Sep 9;6:32819.

68. Hartmannová H, Piherová L, Tauchmannová K, et al. Acadian variant of Fanconi syndrome is caused by mitochondrial respiratory chain complex I deficiency due to a non-coding mutation in complex I assembly factor NDUFAF6. Hum Mol Genet. 2016 Jul 27.

69. Ellingford JM, Barton S, Bhaskar S, et al. Whole Genome Sequencing Increases Molecular Diagnostic Yield Compared with Current Diagnostic Testing for Inherited Retinal Disease. Ophthalmology. 2016 May;123(5):1143-50.

70. Smedley D, Schubach M, Jacobsen JO, et al. A Whole-Genome Analysis Framework for Effective Identification of Pathogenic Regulatory Variants in Mendelian Disease. Am J Hum Genet. 2016 Sep 1;99(3):595-606.

71. Noll AC, et al. Clinical detection of deletion structural variants in whole-genome sequences. Npj Genomic Medicine 2016 Jul 3.

72. Shekelle PG, Woolf SH, Eccles M, Grimshaw J. Developing clinical guidelines. West J Med. 1999;170(6):348-51.

73. National Academies of Sciences, Engineering, and Medicine, Health and Medicine Division, Board on Health Care Services, Board on the Health of Select Populations, Committee on the Evidence Base for Genetic Testing. An Evidence Framework for Genetic Testing. Washington (DC): National Academies Press (US); 2017 Mar 27.

